# Examining the potential of growing frafra potatoes in polypropylene sacs as a food security strategy in the Bolgatanga Municipality of the Upper East Region, Ghana

**DOI:** 10.64898/2026.09.24.754242

**Authors:** Abonuusum Ayimbire, James Anaba Akolgo, Joseph Akolgo Asampana, Ayisoba Awineseligo, Gabriel Asumboya, Richard Akazotiyele

## Abstract

Populations are denser in the urban areas where there is inadequate arable land for farming. This calls for innovative use of limited spaces in sac farming to produce food to augment household nutrition. The study determined the feasibility of cultivating and comparing the performance of different frafra potato varieties in polypropylene sacs. A Randomized Complete Block Design (RCBD) with six treatments, each having three replications, was used. Each treatment was a variety of frafra potato, cultivated on a polypropylene sac filled with nutrient-enriched soil. The data collected was analyzed using STATA 15.0. The results show that plants having leaves with the largest leaf area in 2021 (20.54 (8.41) cm^2^) and in 2022 (19.98 (7.83) cm^2^) were in the MAL variety. The greatest number of leaves (417.11 (255.26)) were produced by the WAP variety in 2021 but by the MAM variety (349.33 (154.63)) in 2022. Plants having the longest vines in 2021 were in the MAM strain (41.24 (22.01) cm) but in the NAT variety (39.93 (21.54) cm) in 2022. In 2021, the highest yield of tubers was 5.56 (0.23) t/ha in the MAL variety with the lowest yield (1.59 (1.27)) t/ha recorded in the LOV strain. In 2022, the NUP variety produced the highest yield (1.91 (0.40)) (t/ha) with the WAP strain producing the lowest (0.87 (0.50)) (t/ha). The data shows that it is feasible to cultivate frafra potatoes on polypropylene sacs. In the main, all the frafra potato varieties produced considerable quantities of vegetative organ biomass and root tubers with the MAL, NAT and NUP varieties performing better. Therefore, though further investigation is required to draw an appropriate conclusion, these varieties may be recommended for cultivation on polypropylene sacs to produce food for both humans and livestock consumption.

## 1.00 Introduction

The crop, *Solenostemon rotundifolius*, is commonly cultivated in the tropical regions of Africa and Asia. In Ghana, the crop, which is referred to as frafra potato, performs well in the Guinea-Savannah vegetational zone, stretching from the three Northern Regions through Upper East to Upper West Region [4, 9, 20, 23]. The cultivation of the crop is limited to indigenous subsistence farmers. However, it is not a fastidious crop, yields in soils of average nutrient content and has great potential for commercialization. It is resilient, able to survive many days of drought and the associated high temperatures. At maturity, provided there are no rains, the tubers can remain on the farm for as long as the soil is dry and be harvested at the convenience of the farmer [12, 14, 17]. All these features of the crop suggest that it can be farmed on a large scale.

*S. rotundifoilus* is a versatile crop whose vegetative organs are useful as animal feed while the tubers, which are rich in nutrients, vitamins, phytochemicals and minerals, are consumed by humans. It has been found to contain a range of 1.35 (0.04) to 4.81% protein, 18.92 (0.72) to 83.39% carbohydrates, 0.73 to 3.06 (0.05) % crude fiber as well as 0.37 (0.03) to 0.63 (0.04)% of fat. The tubers produced by the crop also contain total phenols of about 82.84 mgGAE/100g, total flavonoids 25.16 mgGAE/100g, as well as free radical scavenging capability of 48.09% [1, 6, 18]. It is no wonder that the crop is reported to have some nutraceutical properties, for which reason it has been named ‘*Coleus dysentericus*’ to depict its ability to help in the treatment of dysentery. Given the nutritive and antioxidant content, the root tubers of the crop have been reported to possess potential use in industry, for example, in bread making [23]. Thus, the frafra potato as a crop has so much nutrient fortification and health benefits that it should not be neglected and limited to traditional peasant farming.

However, in spite of these valuable attributes, the popularity of the frafra potato is steadily declining, with its cultivation being limited to indigenous traditional peasant farmers who lack any knowledge of modern agricultural practices. There is little research focus on the crop, and improved varieties are rarely produced and used by the farmers. The small sizes of the tubers, which make it difficult to achieve appreciable tonnage as well as the inconvenience of eating the same, contribute to its neglect and decline in production [4, 12, 14, 17, 23]. There is, therefore, a need to reverse this trend, given the invaluable attributes of the crop by researching into the diverse varieties of frafra potato to produce and select those that improve the sizes of the root tubers as well as the general yield per hectare. It is also necessary to raise public awareness of the crop’s attributes to increase its market potential, thereby encouraging enlightened people to cultivate it. Novel ways of propagating the crop, such as the use of polypropylene sacs, will not only facilitate its cultivation on limited spaces, as prevails in the cities where populations are dense and more food is required, but also contribute to the fight against hunger in the world.

The Ghana Statistical Service (GSS) estimated that about 23.4% of the citizenry are unable to live on $2.00 daily, which is the international poverty line [11]. Again, estimates from the Comprehensive Food Security and Vulnerability Analysis (CFSVA) revealed that about 12% (3.6 million/30 million) of people in Ghana are food insecure, with those in the cities experiencing the worst conditions [25]. The effects of global warming due to the ever-changing climate, on agriculture have also reduced food production. The result is that the average person living in the developing nations of the world is unable to access sufficient quality food [10]. Cultivating root tuber crops like frafra and sweet potatoes on polypropylene, biodegradable sacs [5] will enhance food production close to households, which meets the climate-smart agriculture principles and improves global food security [15]. Cultivating crops on polypropylene, biodegradable sacs, has the added advantage of utilizing small spaces within households to produce food to augment family nutrition. The practice eliminates the cost of transporting food from the rural areas with poor road networks, processing and packaging food, thereby reducing food prices [21, 22]. The current work was thus designed not only to test the feasibility of cropping frafra potatoes on polypropylene sacs, but also to determine which of the six varieties would produce more vegetative organs, which have potential use as animal feed and/or yield more root tubers for human consumption.

## 2.00 Materials and methods

### 2.01 The Study Area

The research was conducted in a community called Yorogo Gaabiisi (10.8280153N, 0.9380438W), a suburb of Bolgatanga Municipality (10.7875°N, 0.8580°W) in the Upper East Region of Ghana (S1 fig). The frafra potatoes were cultivated in Dukomah Garden simultaneously with sweet potatoes between 25th June and 29th October, 2021. The repeat experiment was between 23rd July and 30th November, 2022 [5]. The objective of the research was to determine the feasibility of cultivating frafra potatoes in polypropylene sacs and to compare the growth and yield potential of the six frafra potato varieties. According to the 2021 population and housing census, out of a total population of 139,864 in the Bolgatanga Municipality, females consisted of 52.38%, as against 47.62% males. Also, 63.82% of them live in the towns, with only 36.16% staying in the rural areas, where farming is the main occupation. The total land area of the Municipality is 326.1 km², giving a population density of 428.9/km² and an annual growth of 2.9% (GSS, 2021). [5] reported that prior to the census, of the 4,668 people in Yorogo, females constituted the majority (50.64%), whereas in Gaabiisi, 50.50% of the 697 people were males.

### 2.02 The design of the experiment

The experiment was conducted in a Randomized Complete Block Design (RCBD) format. There were six treatments, and every treatment had three replications, the same as [5] reported for sweet potatoes. Each treatment was a variety of frafra potato and was cultivated on a polypropylene sac filled with nutrient-enriched, mixed garden soil. Given that the total number of frafra potato varieties was six (6), there were six (6) polypropylene sacs of the same size in the experiment. S1 Table shows how the polypropylene sacs were numbered and the respective variety of frafra potato that was cultivated on each.

**Table 1:** Sac number and the respective variety of frafra potato that was cultivated on it.

| Sac number | Sac 1 | Sac 2 | Sac 3 | Sac 4 | Sac 5 | Sac 6 |
| --- | --- | --- | --- | --- | --- | --- |
| Variety of | Manga moya | Naachem | Maa-Lana | WAAP | Nutsuga | local variety |
| Frafra | (MAM) | Tiir (NAT) | (MAL) | Peisa | peisa (NUP) | (LOV) |
| potato |  |  |  | (WAP) |  |  |

The polypropylene sacs were each divided into three (3) equally spaced rows around it, using a marker, to give three replicate rows per sac. Then, on each row, six equally spaced holes were made as the frafra potato vine stands. The polypropylene sacs were filled with nutrient-enriched mixed soil and placed to stand vertically erect. The distance between any pair of holes (vine stands) in each row was 25.50 cm, whereas that between each pair of adjacent rows was 40.00 cm. Each polypropylene sac had six plant stands on each of the three rows on it. Thus, a total of eighteen (3×6 = 18) of the same variety of frafra potato vines were planted per sac. Given that there were six (6) treatments (sacs), a total of one hundred and eight (6×18 = 108) plants were expected in the experiment. The vine stands were made on each sac by use of a heated metal pipe (S1 Plate).

### 2.03 How the soil was prepared for use in filling the polypropylene sacs [5]

The soil that was used to fill the polypropylene sacs was mixed with organic manure from an abandoned fish pond to improve its nutrient content, as well as produce a sandy loamy soil that has good moisture retention capacity. This enhances tuber formation. The soil was prepared by measuring ten (10) head pans of the normal garden soil and thoroughly mixing it with two (2) head pans of the natural organic manure to form a uniform sandy loamy soil (S2 Plate). The mixed, nutrient-enriched soil was then used to fill the polypropylene sacs. Hence, twelve (12) head pans of the nutrient-enriched mixed soil were used to fill each sac for each treatment. In order to ensure adequate and uniform watering of the soil in the sacs, the soil was intermittently watered during the process of filling.

### 2.04 Sampling and testing of the soil [5]

Samples of the various soils (natural manure, garden and mixed soils) were analyzed to determine their physicochemical properties as well as their capacity to support plant growth [16] prior to planting the frafra potato vines. The mixed soils were sampled after mixing each heap of soil into a polypropylene sac. Each sample was taken by using a hand probe to fetch three subsamples from different parts of the mixed soil in a triangular pattern. The three subsamples collected from each sac were carefully mixed to constitute a composite sample from which two grams (2 g) were then fetched into a black plastic bag and duly labelled. Therefore, in all, six (6) composite samples of the mixed soil were made. Three subsamples each were also taken from the garden soil and natural manure, which were rendered into two composite samples (one each). The total of eight (8) composite soil samples were carried to the Soil Science Laboratory of Kwame Nkrumah University of Science and Technology for analysis. These processes of soil sampling and testing were repeated after the frafra potatoes were harvested to determine the impact of the crops on the soil nutrients.

### 2.05 Sources of the vines

The six varieties of frafra potato vines were collected from Dukomah Garden located in Yorogo Gaabiisi (10.8280153N, 0.9380438W), a suburb of Bolgatanga Municipality (10.7875°N, 0.8580°W), where the experiment was carried out.

### 2.06 Planting

Having filled all the sacs with the mixed soil, they were each watered before the frafra potato vines were planted on them using gloves. Planting was done by inserting a stick into the stand to make a hole of about 8 cm. The vine was then inserted 5 cm into the hole, with the soil being gently pushed in to hold the vine. When planting was completed, all the sacs were again watered. The practice of watering the sacs in the mornings and evenings continued for eleven days to supplement rainfall [8].

### 2.07 Data collection

Data was collected in two-week intervals, beginning from the day of planting. In every treatment, a plant was randomly selected from each of the three repeat rows for data collection. The plant thus selected was tagged by a green tag loosely tied to one of its branches. Hence, each variety or treatment had one plant per row to make up three plants per treatment, tagged for data collection. Altogether, the experiment had a total of eighteen (18) plants from which data was collected (3 x 6 = 18). The data was measured from the following parts of the vines of the frafra potatoes: number and length of leaves (cm), leaf area (cm^2^), length of leaf petiole (cm), number and length of vine branches (cm), number and mass of root tubers (Kg).

### 2.08 Estimation of leaf area (cm^2^)

A leaf of the tagged plant was spread on a graph sheet (without plucking it from the vine) and its outline drawn by use of a pencil (S3 Plate). To estimate the leaf area, A, the number of one-centimeter squares that were:

1. Fully covered by the leaf outline were counted and recorded as B,
2. Half-covered by the leaf outline were counted as C and divided by two to record 0.5C,
3. At least three-quarters covered by the leaf outline were counted as being fully covered and recorded as B_1_
4. At most one-quarter covered by the leaf outline was ignored to compensate for the one-centimeter squares that were at least three-quarters covered by the leaf outline and were assumed to be fully covered and recorded as B_1_.

The leaf area, A, in square centimeters (cm^2^) was then calculated as A = B + B_1_ + 0.5C [4, 5].

### 2.09 Leaf length (cm)

Using a centimeter rule, the length of a leaf of the tagged plant was measured from the tip of the leaf to the part joining the petiole.

### 2.10 Petiole length (cm)

The petiole of a leaf of the tagged plant was measured using a centimeter rule from the beginning of the blade to the part attached to the vine and recorded.

### 2.11 Number of leaves

The number of leaves produced by the tagged plant was counted and recorded.

### 2.12 Vine length (cm)

The length of a vine branch of the tagged plant was measured from the branching point to the end by use of a centimeter rule.

### 2.13 Number of vine branches

The number of vine branches produced by the selected plant was counted and recorded.

### 2.14 Cultural practices

Watering of the frafra potato plants continued every morning and evening for eleven days. This helped the young plants to establish roots in the soil [8]. The plants were again watered whenever there was no rain. Weeding was done manually. The experiment was monitored regularly for signs of pests and disease attack. Pests detected include arthropod larvae, locusts and white flies, which were not of any threat to the frafra potato plants. Despite this, the plants were sprayed with an organic pesticide (bypel) in a dose of 15g per 16L of water, using a knapsack machine.

### 2.15 Number of tubers

During harvesting, the number of root tubers that the plants of each row produced was counted and recorded.

### 2.16 Tuber mass

The mass of root tubers that the frafra potato plants of each row produced was measured and recorded in kilograms (Kg) with the aid of a beam balance.

### 2.17 Plant growth assessment

The growth of the frafra potato plants was assessed by the measurement of the vegetative organs such as the mean number and length of the branches of the vines (cm), the mean length (cm) of the leaf petioles and the number, length (cm) and area of the leaves (cm^2^).

### 2.19 Assessment of the yield of the crop

The mass of root tubers that the frafra potato plants of each row produced, as measured in section 2.16, was used to calculate the total mass of root tubers produced per sac or treatment. To be able to use the relation 2◻rh (◻ = 3.14) to calculate the lateral surface area of the polypropylene sac, the radius, r, as well as the height of the polypropylene sac, h, were measured. The yield of the frafra potatoes in tons per hectare (t/ha) was determined by using the formula

Yield = the mass of tubers produced per sac (tons) / the area of the sac (in hectares).

### 2.20 Data analysis

The raw data sets of the vegetative organs obtained in 2021 and 2022 in Microsoft Excel spreadsheets were imported to STATA 15.0 for analyses and presentation in tables and graphs to enhance clarity of interpretation.

## 3.00 Results and discussion

### 3.01 The physicochemical properties of the mixed soil in the polypropylene sacs

The results of the analyses of the garden soil and natural manure are the same as reported by [5], as the frafra potatoes were cultivated simultaneously with the sweet potatoes, using the same garden soil and natural manure to prepare the mixed soil that was used to fill the polypropylene sacs. Table 2 compares the test results of the garden soil with those of the natural manure. The two types of soil are sandy with the same composition of silt (5.48%), but with sand and clay being slightly different. The garden soil is a little more acidic (pH = 5.38) than the natural manure (pH = 5.89). Nevertheless, the latter is over three (3) times higher in total nitrogen (N), calcium (Ca) and phosphorus (P) content, and more than twelve (12) and five (5) times higher in the content of the trace elements zinc (Zn) and iron (Fe), respectively, than the garden soil. Hence, mixing the natural manure with the garden soil increased the content of nutrients in the mixed soil used to fill the polypropylene sacs.

**Table 2:** Results of analysis of the garden soil and natural manure used in the investigation.

| Parameter tested for | Mean values from testing samples of the garden soil and natural manure |  |
| --- | --- | --- |
|  | Mean values (garden soil) | Mean values (natural manure) |
| **The structure of the soil** |  |  |
| Clay (%) | 2.60 | 2.68 |
| Silt, (%) | 5.48 | 5.48 |
| Sand, (%) | 91.92 | 91.84 |
| Texture class | Sand | Sand |
| **Exchangeable bases** |  |  |
| Ca (cmol/kg) | 2.20 | 7.00 |
| Na (cmol/kg) | 0.12 | 0.16 |
| K (cmol/kg) | 1.80 | 2.33 |
| Mg (cmol/kg) | 1.60 | 3.00 |
| Organic carbon (%) | 0.48 | 1.64 |
| Organic matter (%) | 0.83 | 2.82 |
| pH | 5.38 | 5.89 |
| **Exchangeable acidity** |  |  |
| H (cmol/kg) | 0.458 | 0.454 |
| Al (cmol/kg) | 0.418 | 0.301 |
| Avail. P (mg/kg) | 52.87 | 167.70 |
| Total N (%) | 0.070 | 0.245 |
| **Micronutrients** |  |  |
| Mn, mg/kg | 32.45 | 74.47 |
| Zn, mg/kg | 8.91 | 110.00 |
| Fe, mg/kg | 57.95 | 309.85 |
| Cu, mg/kg | 4.81 | 9.86 |

### 3.02 Results of analyses of the mixed soil used in filling the polypropylene sacs

Table 2 reveals the texture of the mixed soil before planting (BP) and after harvest (AH) to be sandy, having 2.93% of clay and 5.24% of silt BP. The proportion of clay increased to 3.02% while that of the silt decreased to 5.17% AH. However, the proportion of sand reduced slightly from 91.83 % BP to 91.81% AH. Other works showed similar results of soil texture, though the proportions varied from this work. Working in the same Dokumah Garden, [3] reported the content of silt as 1.6% BP and AH and that of clay to be reduced from 15.49% BP to 14.18% AH. According to them, the proportion of sand increased from 82.91% BP to 84.22% AH. In addition, [4] recorded the content of clay to be reduced from 6.36% BP to 5.75% AH, with the proportion of sand also reducing from 84.70% BP to 84.00% AH, compared with the current data. The composition of silt deviated from the current pattern, increasing from 8.94% to 10.30%. The exchangeable cations had concentrations of magnesium (Mg) (2.17), sodium (Na) (0.12), potassium (K) (2.24) and calcium (Ca) (3.27) in centimoles per kilogram (cmol/kg) BP. The concentrations of Mg (2.10) and Na (0.08) were reduced while those of K (3.13) and Ca (3.30) increased AH. The proportion of organic matter (0.88%) and organic carbon (1.51%) BP also reduced to 0.78% and 2.10%, respectively, AH. The data agrees with [3], who recorded an increase in concentration of Ca (1.93-2.2), K (0.231-0.24) and Mg (0.87–1.47) AH and a reduction in the concentration of Na (0.0027 - 0.0022) AH. On the other hand, [4] found only reductions in the concentrations of these ions AH: Ca (3.2 - 2.98), Na (0.05 - 0.03), K (0.65 - 0.31) and Mg (1.65 - 1.03), which was attributed to the plants absorbing and assimilating them. The concentrations being reported are much higher than earlier findings [3, 4] and this may be due to the enrichment of the garden soil with the natural manure. There was a little reduction in the pH of the mixed soil from 4.47 BP to 4.43 AH (Table 3), differing from the findings of [5] on sweet potatoes, where the soil became less acidic AH. This suggests that whereas the sweet potato plants caused a slight decrease in the acidity of the soil, frafra potatoes caused an increase in the soil acidity. The concentration of hydrogen ions (H^+^) BP (0.55 cmol/kg) increased slightly to 0.58 cmol/kg AH, confirming the reduction in acidity of the mixed soil after harvesting. This may corroborate the postulate that different species and varieties of plants have different effects on the acidity of soils [5].

**Table 3:** Results of analysis of the mixed soil before planting (BP) and after harvest (AH).

| Parameter tested for | Mean values from analysis of soil samples |  |
| --- | --- | --- |
|  | Mean values before planting (BP) | Mean values after harvest (AH) |
| **The structure of the soil** |  |  |
| Clay (%) | 2.93 | 3.02 |
| Silt, (%) | 5.24 | 5.17 |
| Sand, (%) | 91.83 | 91.81 |
| Texture class | Sand | Sand |
| **Exchangeable bases** |  |  |
| Ca (cmol/kg) | 3.27 | 3.30 |
| Na (cmol/kg) | 0.12 | 0.08 |
| K (cmol/kg) | 2.24 | 3.13 |
| Mg (cmol/kg) | 2.17 | 2.10 |
| Organic carbon (%) | 0.88 | 0.78 |
| Organic matter (%) | 1.51 | 1.34 |
| pH | 4.47 | 4.43 |
| **Exchangeable acidity** |  |  |
| H (cmol/kg) | 0.55 | 0.58 |
| Al (cmol/kg) | 0.36 | 0.37 |
| Avail. P (mg/kg) | 103.68 | 85.77 |
| Total N (%) | 0.13 | 0.13 |
| **Micronutrients** |  |  |
| Mn, mg/kg | 46.83 | 36.64 |
| Zn, mg/kg | 23.50 | 21.02 |
| Fe, mg/kg | 85.47 | 74.28 |
| Cu, mg/kg | 5.62 | 3.72 |

### 3.03 Growth and yield of the frafra potato varieties

Table 4 presents the results of the measurements of growth and yield of the six varieties of frafra potatoes. On the other hand, Table 5 presents the correlation of these parameters with each other. S4 Plate to S9 Plate show the growth stages of the frafra potato plants from a week old to maturity, whereas S10 Plate to S12 Plate portray the harvested frafra potato tubers, which were used to estimate the yield of the crops.

**Table 4:** The mean values from the measurement of growth and yield parameters of frafra potato varieties.

**a. 2021:**
| FPV | Leaf area<br>(cm<sup>2</sup>) | Petiole<br>length<br>(cm) | Leaf<br>length<br>(cm) | Vine length<br>(cm) | Mean<br>number of<br>leaves | Mean<br>crown<br>width (cm) | Tuber yield<br>(t/h) |
| --- | --- | --- | --- | --- | --- | --- | --- |
| LOV | 16.56<sup>ab1</sup><br>(4.54) | 3.86<sup>a1</sup><br>(0.87) | 4.99<sup>a1</sup><br>(0.80) | 37.82<sup>a1</sup><br>(20.46) | 317.44<sup>a1</sup><br>(117.99) | 32.01<sup>a1</sup><br>(13.69) | 1.59<sup>a1</sup><br>(1.27) |
| MAL | 20.54<sup>a1</sup><br>(8.41) | 3.67<sup>a1</sup><br>(1.23) | 5.49<sup>a1</sup><br>(1.35) | 34.40<sup>a1</sup><br>(19.79) | 277.59<sup>a1</sup><br>(156.41) | 31.30<sup>a1</sup><br>(14.42) | 5.56<sup>bc1</sup><br>(0.23) |
| MAM | 13.84<sup>bb1</sup><br>(5.07) | 3.66<sup>a1</sup><br>(1.23) | 4.67<sup>a1</sup><br>(1.16) | 41.24<sup>a1</sup><br>(22.01) | 370.82<sup>a1</sup><br>(169.20) | 31.01<sup>a1</sup><br>(15.92) | 3.02<sup>bde1</sup><br>(0.83) |
| NAT | 13.81<sup>b1</sup><br>(5.12) | 3.66<sup>a1</sup><br>(1.24) | 4.82<sup>a1</sup><br>(1.14) | 39.72<sup>a1</sup><br>(21.86) | 338.04<sup>a1</sup><br>(182.59) | 33.73<sup>a1</sup><br>(16.60) | 1.91<sup>adfg1</sup><br>(0.79) |
| NUP | 15.40<sup>a1</sup><br>(7.27) | 3.95<sup>a1</sup><br>(1.42) | 5.04<sup>a1</sup><br>(1.53) | 41.08<sup>a1</sup><br>(23.81) | 338.19<sup>a1</sup><br>(158.95) | 32.15<sup>a1</sup><br>(13.92) | 2.54<sup>bdegi1</sup><br>(1.96) |
| WAP | 13.14<sup>a1</sup><br>(5.88) | 3.91<sup>a1</sup><br>(1.53) | 4.84<sup>a1</sup><br>(1.52) | 40.10<sup>a1</sup><br>(25.16) | 417.11<sup>a1</sup><br>(255.26) | 32.32<sup>a1</sup><br>(13.76) | 0.79<sup>adfhj1</sup><br>(0.23) |
| Mean | 15.55<br>(6.60) | 3.79<br>(1.25) | 4.98<br>(1.28) | 39.06<br>(22.04) | 343.20<br>(180.82) | 32.09<br>(14.56) | 2.57 (1.84) |

**b. 2022:**
| FPV | Mean<br>leaf area<br>(cm<sup>2</sup>) | Mean<br>petiole<br>length<br>(cm) | Mean leaf<br>length<br>(cm) | Mean vine<br>length (cm) | Mean<br>number of<br>leaves | Mean<br>crown<br>width (cm) | Mean tuber<br>yield (t/h) |
| --- | --- | --- | --- | --- | --- | --- | --- |
| LOV | 16.31<sup>a1</sup><br>(4.36) | 4.03<sup>a1</sup><br>(0.66) | 5.02<sup>a1</sup><br>(0.63) | 35.49<sup>a1</sup><br>(17.75) | 349.33<sup>a1</sup><br>(154.63) | 31.53<sup>a1</sup><br>(12.50) | 1.11<sup>a1</sup><br>(0.61) |
| MAL | 19.98<sup>ab2</sup><br>(7.83) | 4.34<sup>ab2</sup><br>(0.88) | 5.52<sup>a1</sup><br>(1.00) | 32.95<sup>a1</sup><br>(18.28) | 281.56<sup>a1</sup><br>(144.06) | 30.40<sup>a1</sup><br>(13.67) | 1.11<sup>a2</sup><br>(0.61) |
| MAM | 16.60<sup>ab1</sup><br>(6.30) | 3.34<sup>ac1</sup><br>(1.00) | 5.03<sup>a1</sup><br>(0.98) | 30.78<sup>a1</sup><br>(15.72) | 318.15<sup>a1</sup><br>(201.54) | 30.30<sup>a1</sup><br>(14.13) | 1.75<sup>bde2</sup><br>(0.6) |
| NAT | 13.80<sup>ac1</sup><br>(4.96) | 3.86<sup>ab1</sup><br>(0.79) | 4.97<sup>a1</sup><br>(0.79) | 39.93<sup>a1</sup><br>(21.54) | 328.67<sup>a1</sup><br>(195.62) | 33.79<sup>a1</sup><br>(16.52) | 1.11<sup>acfg2</sup><br>(0.23) |
| NUP | 15.13<sup>ab1</sup><br>(6.70) | 3.63<sup>ab1</sup><br>(1.13) | 4.82<sup>a1</sup><br>(1.24) | 35.61<sup>a1</sup><br>(19.47) | 287.78<sup>a1</sup><br>(169.25) | 29.80<sup>a1</sup><br>(13.00) | 1.91<sup>bdehi1</sup><br>(0.40) |
| WAP | 13.80<sup>ac1</sup><br>(5.46) | 3.42<sup>ac1</sup><br>(1.05) | 4.75<sup>a1</sup><br>(1.11) | 38.76<sup>a1</sup><br>(24.86) | 318.44<sup>a1</sup><br>(167.10) | 27.79<sup>a1</sup><br>(13.62) | 0.87<sup>acfgj1</sup><br>(0.50) |
| Mean | 15.94<br>(6.32) | 3.77<br>(0.98) | 5.02<br>(1.00) | 35.59<br>(19.76) | 313.99<br>(172.27) | 30.60<br>(13.87) | 1.31 (0.63) |

**Table 5:** Correlation of growth and yield parameters.

|  | Leaf area | Leaf length | Petiole length | Vine length | Crown width | Number of leaves | Number of tubers | Mass of tubers | yield |
| --- | --- | --- | --- | --- | --- | --- | --- | --- | --- |
| Leaf area | 1.00 |  |  |  |  |  |  |  |  |
| Leaf length | 0.78 | 1.00 |  |  |  |  |  |  |  |
| Petiole length | 0.72 | 0.89 | 1.00 |  |  |  |  |  |  |
| Vine length | 0.45 | 0.50 | 0.56 | 1.00 |  |  |  |  |  |
| Crown width | 0.62 | 0.65 | 0.71 | 0.72 | 1.00 |  |  |  |  |
| Number of leaves | 0.55 | 0.63 | 0.70 | 0.64 | 0.85 | 1.00 |  |  |  |
| Number of tubers | -0.00 | -0.07 | -0.06 | 0.03 | 0.01 | -0.01 | 1.00 |  |  |
| Mass of tubers | 0.11 | 0.00 | -0.10 | -0.03 | 0.02 | -0.05 | 0.75 | 1.00 |  |
| Yield | 0.11 | 0.00 | -0.10 | -0.03 | 0.02 | -0.05 | 0.75 | 1.00 | 1.00 |

### 3.04 The leaf area of the frafra potatoes

The mean leaf area of the frafra potato plants grown on the polypropylene sacs was 15.55 (6.60) in 2021, which slightly increased to 15.94 (6.32) in 2022. These were much smaller than those reported by [4] of 19.6 (6.8). The largest leaves of the frafra potatoes in 2021, which measured averagely 20.54 (8.41) cm^2^ were of the MAL variety, with the WAP variety having the smallest leaves with a mean leaf area of 13.14 (5.88) cm^2^. In a similar vein, in 2022, the MAL variety again recorded the highest leaf area of 19.98 (7.83) cm^2,^ which was not significantly different from the 2021 figure (P = 0.80). However, the WAP (13.80 (5.46)) cm^2^ and NAT 13.80 (4.96) cm^2^ varieties recorded the smallest leaf area in 2022, which were not significantly different from their respective 2021 figures (P = 0.67) and (P = 1.00). In 2021, the leaf area of the MAL variety was significantly larger than that of MAM (P = 0.00), NAT (P = 0.00) and NUP (P = 0.04) varieties only. The leaf areas of the rest of the varieties did not differ statistically from each other. A similar trend was recorded in 2022, where the leaf area of MAL was only significantly higher than that of NAT (P = 0.00) and WAP (P = 0.00). [4] reported a leaf area of 21.2 (7) cm^2^, produced by the LOV variety as the highest and 17.4 cm^2^ recorded in the MAM one as the lowest. These contrast differently with the current data, as MAL provided the largest leaves in both 2021 and 2022. However, MAM produced the smallest leaf area in both studies. Again, the current record of 6.60 and 6.32 standard deviations compares favourably with their value of 6.80, confirming a wide spread of leaf area across the frafra potato plant varieties. Physiologically, leaf area has a proportional relationship with water loss through transpiration and evaporation. Hence, the plants with the largest leaves, like MAL in the current study, may require more watering, especially when cultivated on polypropylene sacs to compensate for the water loss.

Table 5 shows a weak positive relationship between the leaf area and both mass and yield of tubers (r = 0.11). The table also portrays a range of strong positive correlations of leaf area to the mean values of the vegetative organs (r = 0.45 - 0.78). This may account for the phenomenon of plants usually using all the food synthesized by the leaves to produce vegetative organs until maturity or when there is a stressor from the environment to trigger the reproductive phase to commence flowering and fruiting or tuber formation [3, 4, 5, 24]. This may also explain reports by various workers that if the sizes of the leaves of a plant increased to a certain threshold, there would be a natural tendency for the plant to channel its photosynthesized food to the production of vegetative organs at the expense of yield [3, 4, 5, 13, 19, 24]. It may further account for the negative relationship between the current leaf area and the number of tubers (r = −0.0018) because the small sizes of the frafra potato tubers imply that high numbers were produced, which is not correlating favourably with leaf area [3, 4, 5, 24].

### 3.05 The petiole length of the frafra potatoes

The mean petiole length of the frafra potatoes in 2021 was 3.79 (1.25) cm. The longest (3.95 (1.42) cm) and shortest petioles (3.66 (1.23) cm) were produced by the NUP and MAM varieties, respectively. In 2022 on the other hand, the MAL variety recorded the longest leaf stalk (4.34 (0.88) cm) with the MAM strain again producing the shortest ones (3.34 (1.00) cm), and the difference between the two was significant (P = 0.01). The leaf stalks of the MAM variety were also significantly longer than those of the WAP strain (P = 0.03). However, there were no significant differences in leaf stalk length across the rest of the varieties. The length of the leaf stalks is positively related to the other vegetative organs (vine length (r = 0.56), crown width (r = 0.71), number of leaves (r = 0.70) but negatively to the number of tubers (r = −0.06), mass of tubers (r = −0.10) and yield (r = −0.10).

### 3.06 The leaf length of the frafra potatoes

During the 2021 crop season, the frafra potato plants produced a mean leaf length of 4.98 (1.28) cm with no significant variations across the varieties (0.28≤P≥1.00). The longest leaves were recorded by the MAL variety (5.49a1 (1.35) cm), while the shortest leaves were produced by the MAM strain (4.67a1 (1.16) cm). Whereas the MAL variety again produced the longest leaves in 2022, the shortest leaves were found in the WAP strain (4.75 (1.11) cm) with no statistical difference across the varieties (0.07≤P≥1.00). Leaf length correlated strongly with the other vegetative organs; leaf area (r = 0.78), petiole length (r = 0.89), vine length (r = 0.50), crown width (r = 0.65) and number of leaves (r = 0.63). It, however, correlated weakly with the mass (r = 0.00) and yield (r = 0.00) of tubers but negatively with the number of tubers (r = −0.07). This is in agreement with the observation that plants first produce vegetative organs before tubers are formed later.

### 3.07 The vine length of the frafra potatoes

The longest frafra potato vines were recorded in the MAM variety in 2021 (41.24 (22.01) cm) but in the NAT (39.93 (21.54) cm) strain in 2022, although both were not statistically different from each other in length (P = 0.84). With a mean length of 39.06 (22.04) cm in 2021 and 35.59 (19.76) cm in 2022, the vines of the different frafra potato varieties were not statistically longer than each other in both years (P=1.00). The shortest vine branches were observed in MAL 34.40 (19.79) in 2021 but in the MAM 30.78a1 (15.72) in 2022. The data reveals vine length to be strongly correlated to crown width (r = 0.72), number of leaves (r = 0.64), vine length (r = 0.45) but weakly correlated to number of tubers (r = 0.03) and negatively related to mass (r = −0.03) and yield (r = −0.03) of tubers (Table 5). Thus, naturally, frafra potato vines grow at the expense of tuber formation and hence, yield.

### 3.08 The number of leaves produced by the frafra potato plants

The plants produced a mean number of leaves of 343.20 (180.82) in 2021 and 313.99 (172.27) in 2022. In 2021, the MAL variety recorded the least number of leaves (277.59 (156.41)) as the WAP strain produced the greatest number of leaves (417.11 (255.26)). In consonance with the 2021 data, the MAL variety again had the least number of leaves of 281.56a1 (144.06) in 2022, while the MAM strain produced the highest number of leaves of 349.33 (154.63). There was no statistical difference in leaf number across the six varieties in both years (0.07≤P≥1.00). The current work reveals that varieties having small leaf areas tend to have a large number of leaves to compensate for the small leaf sizes (Table 4). Thus, in 2021, the MAL variety with the largest leaf area of 19.98 (7.83) cm^2^ recorded the smallest number of leaves (281.56 (144.06)), and in 2022, the same strain produced the widest leaves of mean area 19.98 (7.83) cm^2^ and also recorded the least number of leaves (281.56 (144.06). On the other hand, in 2021, the WAP variety, which had the smallest leaf area (13.14 (5.88) cm^2^), produced the largest number of leaves of 417.11 (255.26). In 2022, the NAT strain having the least leaf area, 13.80 (4.96) cm^2^, recorded the second largest number of leaves of 328.67 (195.62). This is because physiologically, photosynthesis is directly related to leaf area so that small leaves have reduced photosynthesis and ought to offset this by increasing the number of leaves.

The number of leaves, followed the same trend of strong correlation with the other vegetative organs, ranging from leaf area (r = 0.56) to crown width (r = 0.85) but with negative correlation to the number (r = −0.01), mass (r = −0.05) and yield of tubers (r = −0.05), reflecting the interdependence of the vegetative organs on each other and a tendency to grow at the expense of tuber formation and hence, yield.

### 3.09 The crown width of the frafra potatoes

The widest (33.73 (16.60) cm) and narrowest (31.01 (15.92) cm) crowns in 2021 were observed in the NAT and MAM varieties, respectively, though these were not statistically different from each other (P = 1.00). The NAT variety again had the widest crowns (33.79 (16.52) cm) with the WAP one recording the narrowest crowns (27.79 (13.62)) in 2022, which were also not significantly wider than each other (P = 1.00). Crown width correlated positively with leaf area (r = 0.62) and vine length (r = 0.72). In terms of biomass production, the MAL, MAM and NAT varieties were the best performers in both years (Table 4). The mean crown width was 32.09 (14.56) in 2021 and 30.60 (13.87) in 2022, which were both wider than the 27.9 (90) cm reported by [4]. A high positive correlation of crown width to leaf area (r = 0.78 – 0.94) was interpreted to reflect their dependence on each other [4]. In the current results, the relationship of crown width to the other vegetative organs ranges from leaf area (r = 0.62) to the number of leaves (r = 0.85), again reflecting the interdependence of the vegetative organs on each other. There was, however, a weak correlation with the number (r = 0.01) and mass (r = 0.02) of tubers as well as yield of tubers (r = 0.02), again indicating the natural tendency of the vegetative organs to thrive at the expense of tuber formation and hence, yield.

### 3.10 The root tuber yield of the frafra potatoes

The mean tuber yield in 2021 was 2.57 (1.84) tons per hectare (t/ha), which was reduced to 1.31 (0.63) (t/ha) in 2022. In 2021, the highest yield of frafra potato tubers was 5.56 (0.23) in the MAL variety, which was significantly higher than the lowest yield of 0.79 (0.23) in the WAP strain (P = 0.00). The LOV variety produced 1.59 (1.27) t/ha, and this was significantly lower than the yield of MAL (5.56 (0.23) t/ha), MAM (3.02 (0.83) t/ha), P = 0.00 and NUP (2.54 (1.96) t/ha), P = 0.02.

In 2022, the NUP variety produced the highest yield of 1.91 (0.40) (t/ha) with WAP producing a significantly lower yield of 0.87 (0.50) (t/ha) (P = 0.00). During the 2022 season, the LOV (1.11 (0.61) t/ha), MAL (1.11 (0.61) t/ha), and NAT (1.11 (0.23)) varieties produced significantly lower yield than MAM (1.75 (0.6) t/ha) (P = 0.00) and NUP (1.91 (0.40) t/ha) (P = 0.00). Thus, MAM and NUP were the best yielding varieties in both seasons, whereas MAL was the best yielding variety in only 2021. In general, however, the frafra potatoes cultivated on polypropylene sacs performed better in terms of root tuber yield in 2021 than in 2022.

In contrast, cultivated on ridges under rain-fed conditions, [4] recorded the highest and lowest tuber yields in the LOV (27.8 (4.7) t/ha) and MAM (15.1 (6.4) t/ha) varieties, respectively. Thus, the yield potential of the varieties of frafra potato is not static. The highest yielding variety changed from Lov on ridges [4] to MAL in 2021 and NUP in 2022 on polypropylene sacs, suggesting that yield capacity may not be dependent on just the variety, but other factors may also be influential. Other workers postulated that different varieties respond to various soil conditions in diverse ways, leading to dissimilar results with respect to growth and yield [2, 7]. Again, the yield realized when the same frafra potato varieties were cultivated on ridges was much higher than when grown on polypropylene sacs in 2021, which were also higher than in 2022, implying a need for further assessment to improve yield on the sacs.

### 3.11 Vegetative growth rate of the frafra potato varieties

The leaf area of the plants, expressed in square centimeters (cm^2^), increased continuously irrespective of rainfall in 2021 (S2 fig). This may be attributed to the supplementary watering whenever it did not rain. The leaf area growth of MAL and WAP varieties declined slightly at the 14^th^ and 16^th^ weeks after planting, respectively. The leaf area of the other varieties appeared to continue expanding even at the 18^th^ week after planting. This contrasts with the report of [4] on the same varieties grown on ridges under rain-fed conditions, that the leaf areas stopped expanding when the plants matured. Thus, the current trend might be due to the continuous supply of water to the plants through the supplementary watering. Under rain-fed conditions, the plants experienced the stress of intermittent droughts, which probably triggered the arrest of vegetative growth to flower and form tubers. This may also explain the better mean yield observed under rain-fed conditions of 19.6 (6.8) t/ha [4]. In the present study, in the absence of water stress, the plants continued to use the food made by the leaves to produce vegetative biomass at the expense of tuber formation due to the high correlation between the production of vegetative organs and leaf area (Table 5). The continued increase in leaf area likely exceeded the threshold, for which reason the photosynthates were channeled into the production of biomass at the expense of tuber formation, which explains the better yield under rain-fed conditions [3, 4, 5, 24].

In 2022, the leaf area of the MAM variety dipped from the 10^th^ to the 12^th^ weeks after planting before regaining increasing growth from the 14^th^ week (S3 fig). That of the MAL strain declined from the 14^th^ week and hardly recovered, while the NUP variety took the first four weeks to establish itself before growing steadily to dip slightly at the 16^th^ week. The NAT strain dipped at the 6^th^ week to regain increasing leaf area from the 10^th^ week and remained so to the 18^th^ week after planting. Though it dipped slightly at the 12^th^ week after planting, the WAP variety followed the same trend as in 2021, dipping at the 16^th^ week. The leaves of the frafra potatoes expanded much more in 2022 than in 2021, with some exceeding 500 cm^2^ as against 140 in 2021. This might explain why in 2021 the leaf areas continued to increase with little sign of stagnating, unlike what was observed in the 2022. The high rate of vegetative biomass production in 2022 might also have been at the expense of tuber formation, resulting in the poorer yield.

## 4.00 Conclusion

The data show that it is feasible to grow frafra potatoes on polypropylene sacs for the production of both vegetative organ biomasses to feed livestock as well as for root tubers to feed humans. In the main, all the frafra potato varieties produced considerable quantities of vegetative organ biomass with the MAL and NAT varieties producing the largest quantities while the MAM strain recorded the least. The MAL and NUP strains also produced the highest root tuber yield. Therefore, though further investigations are required to make a conclusion, the MAL, NAT and NUP varieties may be recommended for cultivation on polypropylene sacs for both human food and livestock feed production.

## 5.00 Recommendations

i. Further investigations are needed on the cultivation of these varieties on polypropylene sacs to gather adequate data to recommend which varieties are most suitable.
ii. More investigations should be conducted on the cultivation of other varieties on polypropylene sacs to be able to properly recommend which varieties are most suitable.
iii. Farmer education is required for farmers to accept the innovation and practice sac farming.

## Acknowledgements

We wish to express our profound appreciation to Dukomah Garden for providing the frafra potato varieties for the experiments in both 2021 and 2022. We are also grateful to Mr. Alolga Johnson for providing us with STATA 15.0 and assisting in the data analysis.

## Contributions of Authors

All authors collaborated in contributing resources to conduct the research. Conceptualization, methodology and writing of the original draft were by Author AyA who also supervised Author AwA to conduct the investigation (cultivate the crops) with the help of Author AR. Author AnJA and Author AG performed Data Curation while Author AsJA carried out the formal analysis validation of the data. Review and editing were performed by all the authors who have read and approved the final version of the manuscript.

## Data availability

Data is available in the supporting information files as S1_data analysis.doc and S2_data analysis.doc and S3 data.Exl

## Ethical clearance

The study did involve human subjects. Permission to publish the pictures of Mr. Ayisoba Awineseligo and the map of the study are also submitted.

## Supporting information

S1 Plate: Creation of plant stands on the polypropylene sacs

S2 Plate: The natural manure and garden soil used to fill the polypropylene sacs

S3 Plate: Tracing the outline of a leaf on a graph sheet

S4 Plate: A week-old frafra potatoes cultivated in 2022

S5 Plate: A month-old frafra potatoes cultivated in 2022

S6 Plate: Two-month-old frafra potatoes cultivated in 2022

S7 Plate: Three months of frafra potatoes cultivated in 2021

S8 Plate: Four-month-old frafra potatoes cultivated in 2022

S9 Plate: Frafra potatoes just before harvesting in 2021

S10 Plate: A sample of harvested frafra potatoes from a sac

S11 Plate: Frafra potato tubers per sac, harvested in 2021

S12 Plate: Frafra potato tubers per sac, harvested in 2022

---

**S1 fig:**
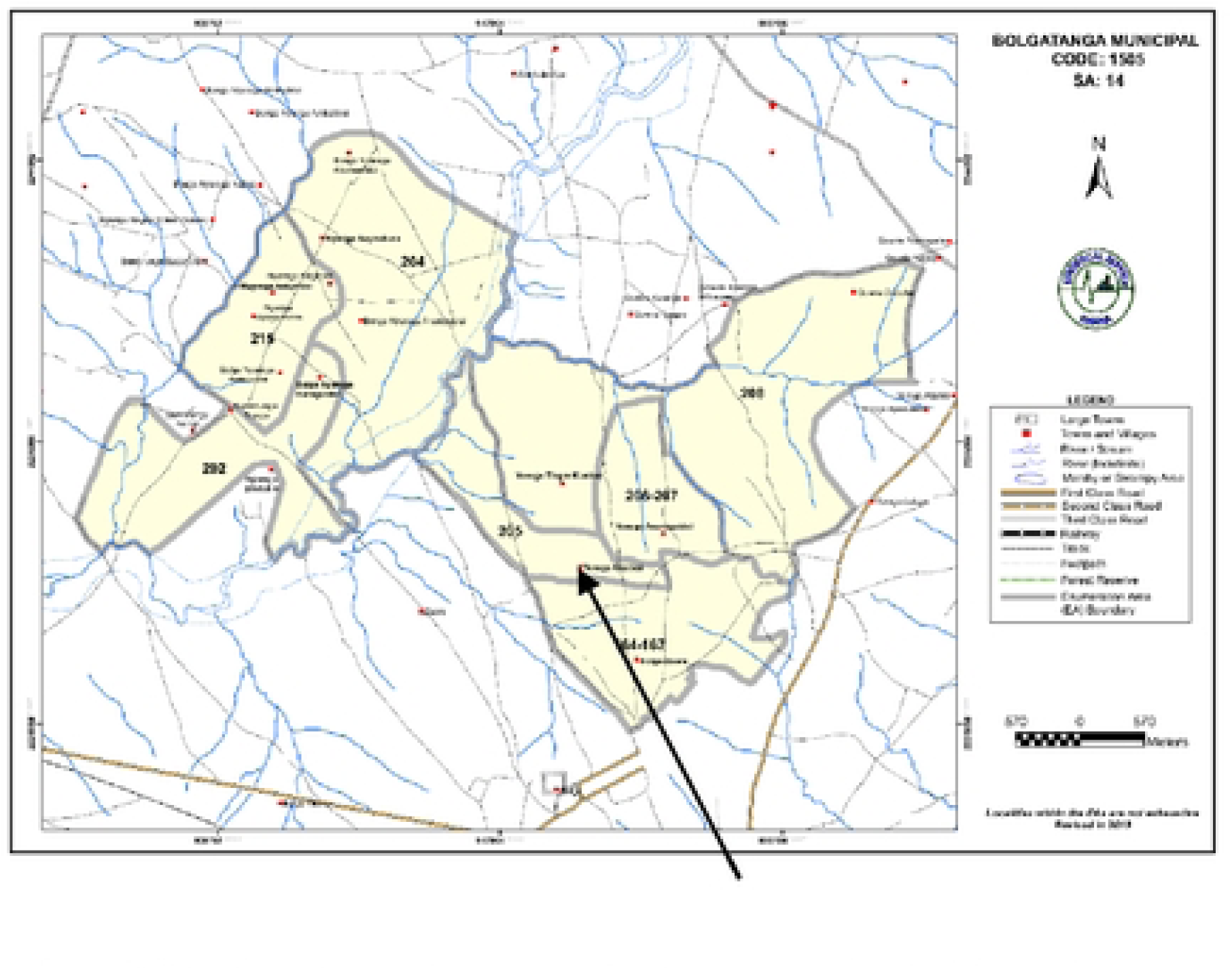
A Map of part of Bolgatanga Municipality. The arrow points at Yorogo Gaabiisi (10.8280153N, 0.9380438W), the study area.

**S2 fig:**
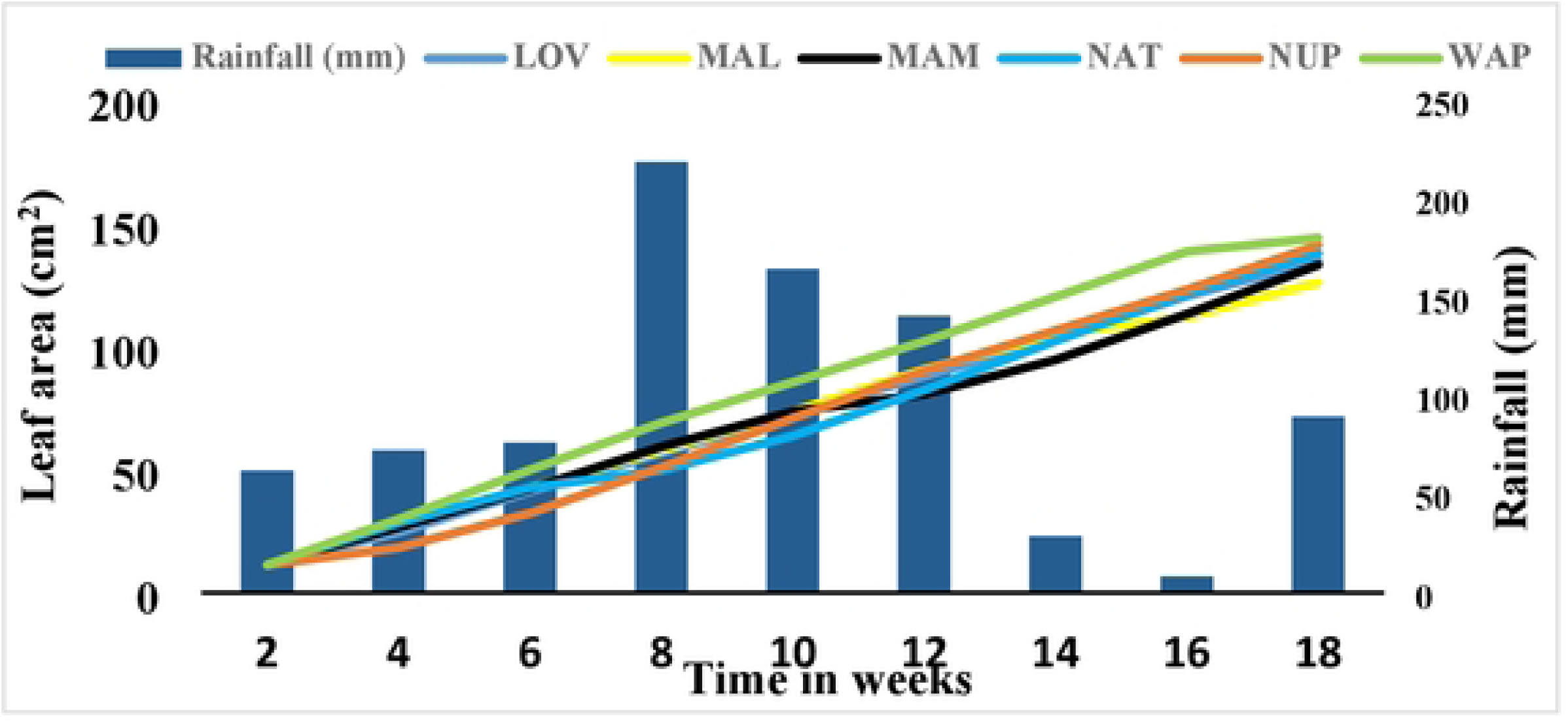
Comparing the rate of increase in leaf area of six frafra potato varieties cultivated on polypropylene sacs in 2021 with time.

**S3 fig:**
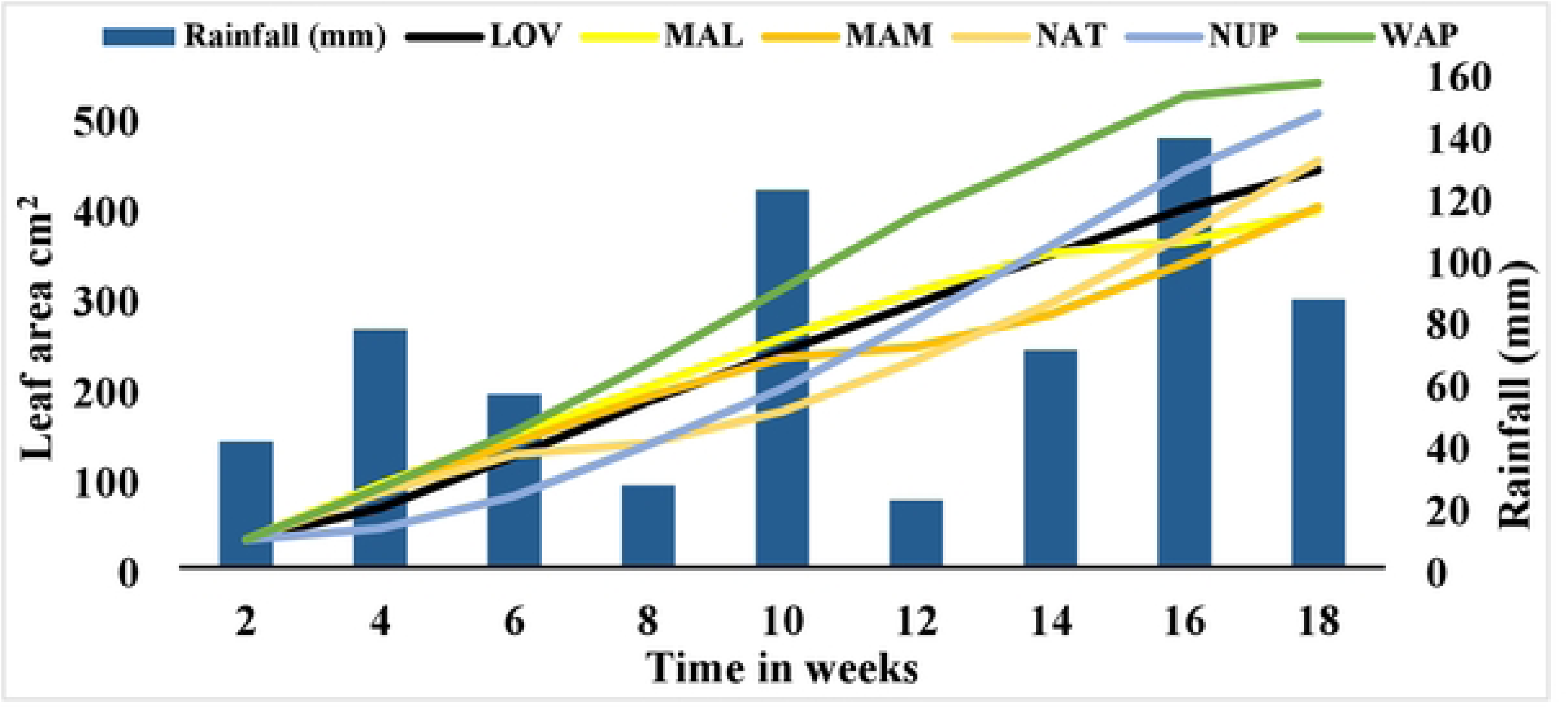
Comparing the rate of increase in leaf area of six frafra potato varieties cultivated on polypropylene sacs in 2022 with time.

